# Rewiring of RNA–protein coupling in osteocytes in response to hyperglycemic levels of glucose

**DOI:** 10.64898/2025.12.25.696466

**Authors:** Aseel Marahleh, Sherif Rashad, Eisuke Kanao, Ziqiu Fan, Fumitoshi Ohori, Hideki Kitaura

## Abstract

Osteocytes are central regulators of skeletal homeostasis, yet how their transcriptome and proteome jointly adapt to metabolic stress is unclear. Here, we combined RNA-sequencing with label-free quantitative proteomics in MLO-Y4 osteocytic cells exposed to hyperglycemic levels of glucose, to interrogate RNA–protein coupling at baseline and under hyperglycemic stress. Transcriptionally, high glucose and mannitol elicited overlapping osmoadaptive transcriptional programs indicative of metabolic remodeling, whereas high glucose alone induced mitochondrial and oxidative phosphorylation downregulation alongside selective activation of bone anabolic and inflammatory pathways. RNA–protein integration revealed moderate coupling at baseline, indicating that mRNA levels capture only part of the proteomic output. High glucose reorganized RNA-protein relationship, sorting genes into four patterns of matched or opposing RNA–protein responses. The four groups were enriched in distinct biological pathways that shaped cellular response to stress exposing significant post-transcriptional and translational control under high glucose levels. These data indicate that osteocytes adapt to stress through program-specific RNA–protein interactions in which post-transcriptional regulation of protein translation reshapes the proteome beyond what is predicted by mRNA levels.

## Introduction

Skeletal homeostasis depends on bone cell’s ability to sense and integrate metabolic and mechanical cues. This coordination is achieved through the interplay of osteoblasts, osteoclasts, and osteocytes, which together execute balanced bone remodeling [1]. Among these, osteocytes serve as the central mechanosensory and endocrine regulators, orchestrating the activity of both osteoblasts and osteoclasts [2-5].

Gene expression is regulated through multiple different molecular levels [6], RNA is transcribed and goes into processing, splicing, turnover, and modifications [7-10]. Thus, post-transcriptional RNA processing influences RNA translation efficiency and protein abundance [11]. Post-translational modifications, protein degradation, and quality control mechanisms determine protein levels leading to modulating cellular behavior [12]. These layers of regulation can amplify, buffer, or completely redirect the biological outcomes predicted from mRNA levels. Therefore, cellular behavior post-transcriptionally emerges from interactions across the transcriptome, proteome, post- and intermediate regulatory levels [13-16].

Studies of osteocyte biology have typically examined single regulatory layers in isolation, so relationships between mRNA abundance and proteomic output remain poorly defined [17-19]. Our previous work showed that high-glucose challenge in primary osteocytes elicits extensive post-transcriptional regulation of a wide range of cellular processes despite only modest changes in transcript levels [20]. Osteocytes are highly glycolytic cells, and they are sensitive to levels of glucose exposure [21,22]. Both elevated and restricted glucose levels have been shown to alter osteocyte function and metabolism [23-25]. During differentiation, osteocytes retain a predominantly glycolytic phenotype but acquire a more flexible metabolic profile as they mature [19,26]. However, our current understanding remains fragmented; transcriptional and proteomic responses to perturbed osteocyte metabolism have been studied independently of each other. How these layers are coordinated, decoupled, or rewired under high glucose remains largely unknown.

To address this gap, we combined RNA sequencing with label-free quantitative proteomics of MLO-Y4 cells to define the multi-layered response of osteocytic cells to high glucose. By integrating these datasets, we assessed RNA–protein coupling and evaluated how this regulatory relationship is remodeled under stress. Our results revealed that at baseline conditions osteocytes show a moderate correlation between transcript abundance and protein expression. High-glucose exposure did not alter the baseline RNA-Protein RANK correlation but unmasked distinct regulatory profiles across coupled and uncoupled RNA-protein relationships. Through a bioinformatics integrative analysis, we identified four distinct regulatory patterns of coupled and uncoupled transcription-translation regulation that govern cell behavior in response to high glucose challenge. This work provides a stepping-stone to move beyond expression data alone and begin identifying the early regulatory layers that shape osteocyte function in metabolic disease and open the door for more detailed mechanistic studies.

## Results

### 1. Pathway enrichment analysis reveals distinct transcriptional programs activated under high glucose stress

Cells were cultured in high glucose (HG) or matching osmolarity D-mannitol (M) for 72 hours, after which total RNA was extracted for RNA-sequencing. Differential gene expression (DGE) analysis revealed that both treatments induced minimal changes in global gene expression (Fig. 1A, B; Fig. 1A Sup). To investigate functional adaptations under high glucose and osmotic stress, we performed gene set enrichment analysis (GSEA). The top significantly enriched pathways unique to each treatment are summarized in Supplementary Tables 1-3. Shared pathways enriched in both HG and M conditions were filtered out from GSEA of M compared to normal glucose (NG) and HG compared to NG (Supplementary Table 1 and 2) and visualized separately (Fig. 1C; Supplementary Table 4).

**Figure 1:**
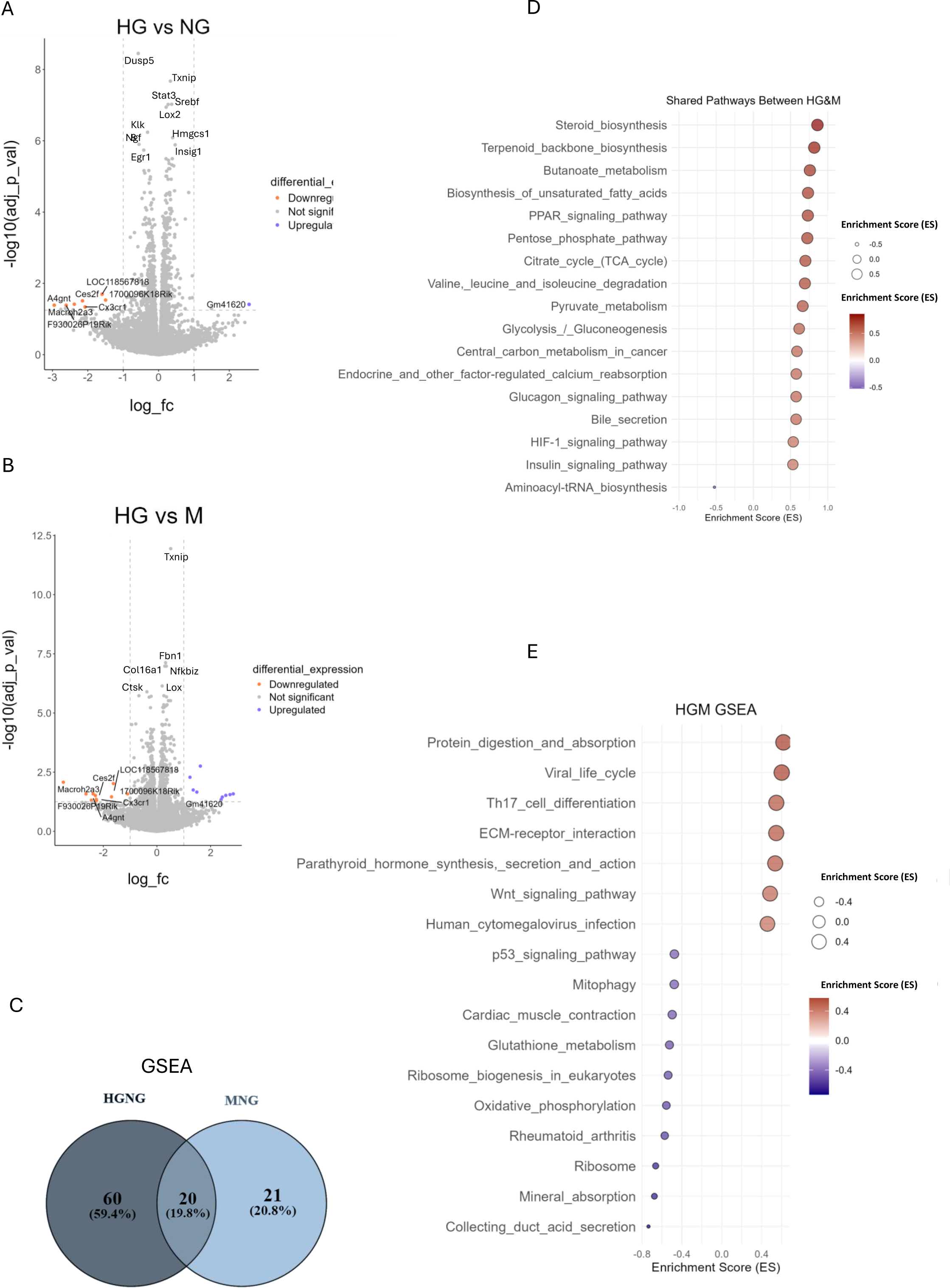
High glucose elicits similar osmodoaptive and glucose-specific transcriptional responses compared to mannitol treatment. **(A)** Volcano plot comparing the log2 FC and - log10 adjusted P value of RNA-seq data in MLO-Y4 cells comparing HG vs NG and **(B)** HG vs M. **(C)** Venn diagram showing the number of shared pathways obtained in GSEA comparing the responses between HGNG and MNG. **(D)** GSEA analysis of pathways shared among HGNG and MNG and between **(E)** M and HG, obtained using the eVITTA toolbox. RNA-seq analysis was done on n=3.

#### 1.1 Pathways common to high glucose and mannitol treatments are enriched in metabolic remodeling

GSEA of cells exposed to HG and M revealed substantial overlaps in the enriched pathways. Prominent among the enriched pathways were those associated with lipid metabolism, including steroid, terpenoid, butanoate, and fatty acid biosynthesis, as well as the peroxisome proliferator-activated receptor (PPAR) signaling pathway featuring *Pparγ* in the leading edge. In parallel, we observed enrichment of metabolic pathways such as the pentose phosphate pathway (PPP), tricarboxylic acid (TCA) cycle, amino acid degradation, and pyruvate and glucose metabolism (Fig. 1D). Pathways involved in ion and solute transport, including calcium ion reabsorption, were also activated. Furthermore, enriched pathways were also consistent with metabolic rewiring such as hypoxia-inducible factor 1 (*Hif1*) and insulin signaling pathways. Notably, both treatments showed reduced enrichment of the aminoacyl-tRNA biosynthesis pathway, suggesting the engagement of a translational control mechanism that suppresses global protein synthesis in response to stress. This similarity likely reflects a shared osmoadaptive response, as both treatments impose comparable high-osmotic stress.

#### 1.2 Distinct high glucose specific pathways reveal oxidative stress and mitochondrial transcriptional enrichment

When osmolarity was controlled for by comparing HG directly to M, distinct transcriptional programs emerged (Fig. 1E; Supplementary Table 3). The top enriched pathways highlighted functional adaptations related to the extracellular matrix (ECM) components, including collagens, proteoglycans, glycoproteins, osteopontin, integrins, and laminins; components essential for maintaining skeletal ECM integrity. In parallel, genes associated with parathyroid hormone (PTH) and PTH-like signaling were significantly enriched. GSEA further revealed activation of inflammatory and immune-related pathways such as Th17 cell differentiation, featuring transforming growth factor receptors (*Tgfbr2*) and several interleukin receptors (*Il6r, Il2r, Il1br*), together with downstream mediators (*Smads, Stats, Mapks, Nfkb*) and growth regulators (Runx2, mTOR, Rar/Rxr). The Wnt signaling pathway was also upregulated, encompassing canonical and non-canonical branches among the leading edge. Interestingly, several pathways annotated as “viral infection” were enriched, containing interferon-like stress response genes commonly associated with innate immune activation under sterile metabolic stress.

Conversely, downregulated pathways converged on oxidative stress responses and ionic balance, including glutathione and mineral metabolism. Among the leading-edge genes, *Mt1, Mt2, Hmox1, Fth1, Ftl1, Atox1, and Atp1a1/b1* indicated suppression of metal ion homeostasis and redox defense mechanisms. A prominent group of downregulated terms was associated with mitochondrial function and oxidative phosphorylation (oxphos), including genes encoding subunits of complexes I–IV of the electron transport chain, ATP synthase (complex V), and proteins involved in mitophagy and H⁺ transport. This pattern extended to vacuolar ATPase components (*Atp6v1, Atp6v0*), which are critical for intracellular pH regulation and bone matrix acidification. Additionally, genes categorized under “cardiac muscle contraction” were suppressed; an annotation encompassing mitochondrial energy production, ion transport (Na⁺/K⁺/Ca²⁺ ATPases and voltage-gated channels), and actin-associated contractile proteins. Of note, the rheumatoid arthritis pathway was also downregulated, with a leading edge including Ctsk and genes involved in monocyte recruitment, osteoclast precursor migration, and osteoclastogenesis. Finally, genes associated with ribosomal biogenesis were also downregulated.

Collectively, these results reveal two distinct adaptive responses to HG. Osmotic stress primarily remodels lipid and glucose metabolism. In contrast, when osmolarity is controlled for, the specific transcriptional effects of HG emerge, dominated by the downregulation of mitochondrial and oxidative phosphorylation pathways, together with reduced expression of genes involved in antioxidant defense.

### 2. Integrative Transcriptome–Proteome Analysis Reveals Layer-Specific Biases at Baseline Conditions

Next, we performed label free proteomics analysis on cells cultured in NG or HG conditions. Our integrative analysis of transcriptome and proteome datasets identified 12623 protein-coding transcripts (Supplementary Table 5) of total transcripts mapped and 4630 transcripts were detected at the protein level, nearly a 36% overlap (Figure 2A, Supplementary Table 6). To examine the relationship between transcript abundance and protein abundance, we generated a joint density plot combining RNA-seq and proteomics datasets from co-expressed and exclusively detected products (Figure 2. B). The central two-dimensional (2-D) histogram represents the distribution of genes detected at both transcript and protein levels, with color intensity corresponding to log₁₀-transformed mean expression values of co-expressed products. The diagonal trend indicates a general positive association between transcript and protein abundance at baseline conditions (Figure 2B).

**Figure 2:**
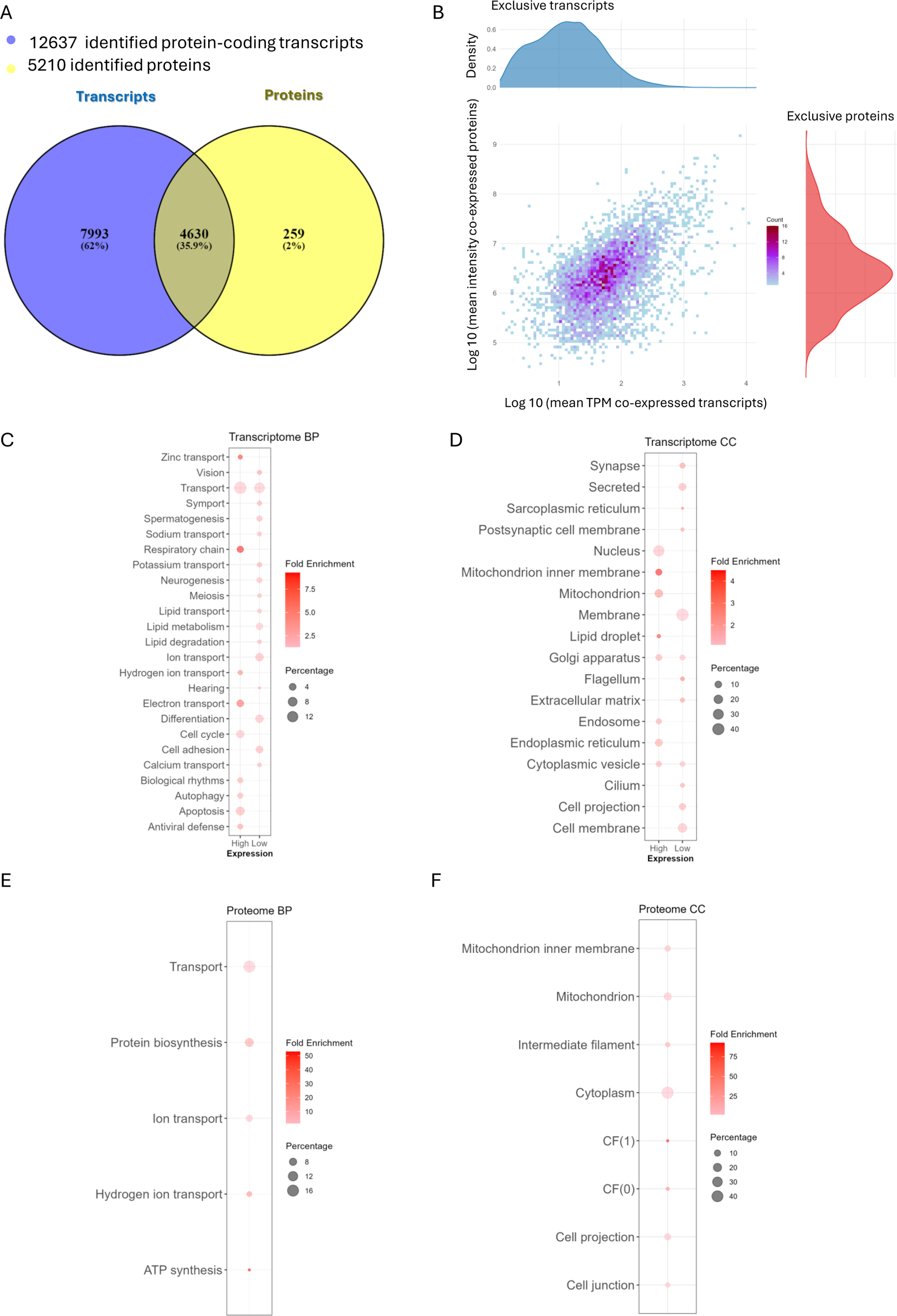
RNA-protein integration shows a moderate correlation at baseline culture conditions. **(A)** Venn diagram shows the number protein coding RNAs, the number of proteins identified, and percentage of products co-expressed at both transcriptomic and proteomics layers. **(B)** Central scatter plot of co-expressed products. Marginal top/blue density plot represents genes exclusively found in RNA-seq, and marginal right/red plot represents genes exclusively found in proteomics analysis. **(C)** GOBP and **(D)** GOCC analysis of transcriptome-exclusive genes. For functional annotation of transcript-exclusive genes, genes were divided into mean log10 TPM values less than 0 and considered lowly expressed, whereas values more than 0 were considered highly expressed. **(E)** GOBP and **(F)** GOCC analysis of proteome-exclusive genes analyzed using the DAVID platform. Percentage represents the percentage of genes found in our datasets relative to genes corresponding to the annotated term. BP= biological process, CC = cellular component.

The marginal density plot above the main 2-D histogram depicts the expression distribution of transcripts detected exclusively at the RNA level, while the right marginal plot shows proteins detected exclusively at the proteomic level. Both marginals align with the respective axes of the co-expression map, allowing direct comparison of expression ranges between exclusive and shared features. The transcript-exclusive density (top) is skewed toward lower log₁₀(TPM) values, suggesting many undetected proteins are likely lowly expressed at the transcriptional level and below the proteomic detection threshold, whereas the protein-exclusive distribution (right) occupies a narrower, moderate-intensity range (Figure 2B).

Gene ontology (GO) enrichment analysis was performed using the DAVID platform [27]. Transcripts expressed exclusively at the RNA level were enriched in biological processes (BP), including ion transport, cell adhesion, differentiation, and general transport pathways. Corresponding cellular component (CC) terms were significantly enriched for synaptic regions, secreted and membrane-associated terms, indicating that many transcript-only products are localized to membrane-bound compartments. Together, these findings imply that the exclusive transcriptome predominantly represents mRNAs involved in secretory proteins or are part of a membrane structure. The proteins encoded by these genes are difficult to identify by MS due to their biological characteristics [28], physicochemical properties or specific cellular localization owing to MS inherent technical limitations [29,30] (Figure 2C, D).

GO analysis of products detected exclusively at the proteomic level was dominated by protein biosynthesis, ATP synthesis, and hydrogen ion transport. Corresponding CC categories included the cytoplasm, mitochondrial inner membrane, and ATP synthase complexes. The proteins detected exclusively at the proteomic level were predominantly mitochondrial enzymes and cytoskeletal components. These species were moderately expressed within the proteome but lacked detectable transcripts above the RNA-seq threshold. Many of these proteins, including mitochondrial oxphos subunits, are known to persist independently of acute mRNA levels through slow turnover and efficient recycling (Figure 2E, F) [31]. Their mid-range abundance likely reflects a combination of long protein half-lives, high basal stability, and limited mRNA detectability under the chosen filtering criteria rather than true transcriptional silencing [32,33].

Consistent with this interpretation, the co-expressed set of genes showed a moderate positive correlation between transcript and protein abundance at NG and HG indicating the overall coupling level between the two expression layers (Figure 3A, B). To assess the effect of HG stress on the relationship between mRNA and protein expression changes, we compared the log₁₀ fold changes of co-expressed transcripts and their corresponding proteins. Exposure to HG produced an uncoupled phenotype in our cells (Spearman’s ρ = 0.03, p = 0.019), indicating a dissociation between RNA and protein responses. This uncoupling suggests that HG impacts post-transcriptional regulation of RNA altering translation or post-translational controls.

**Figure 3:**
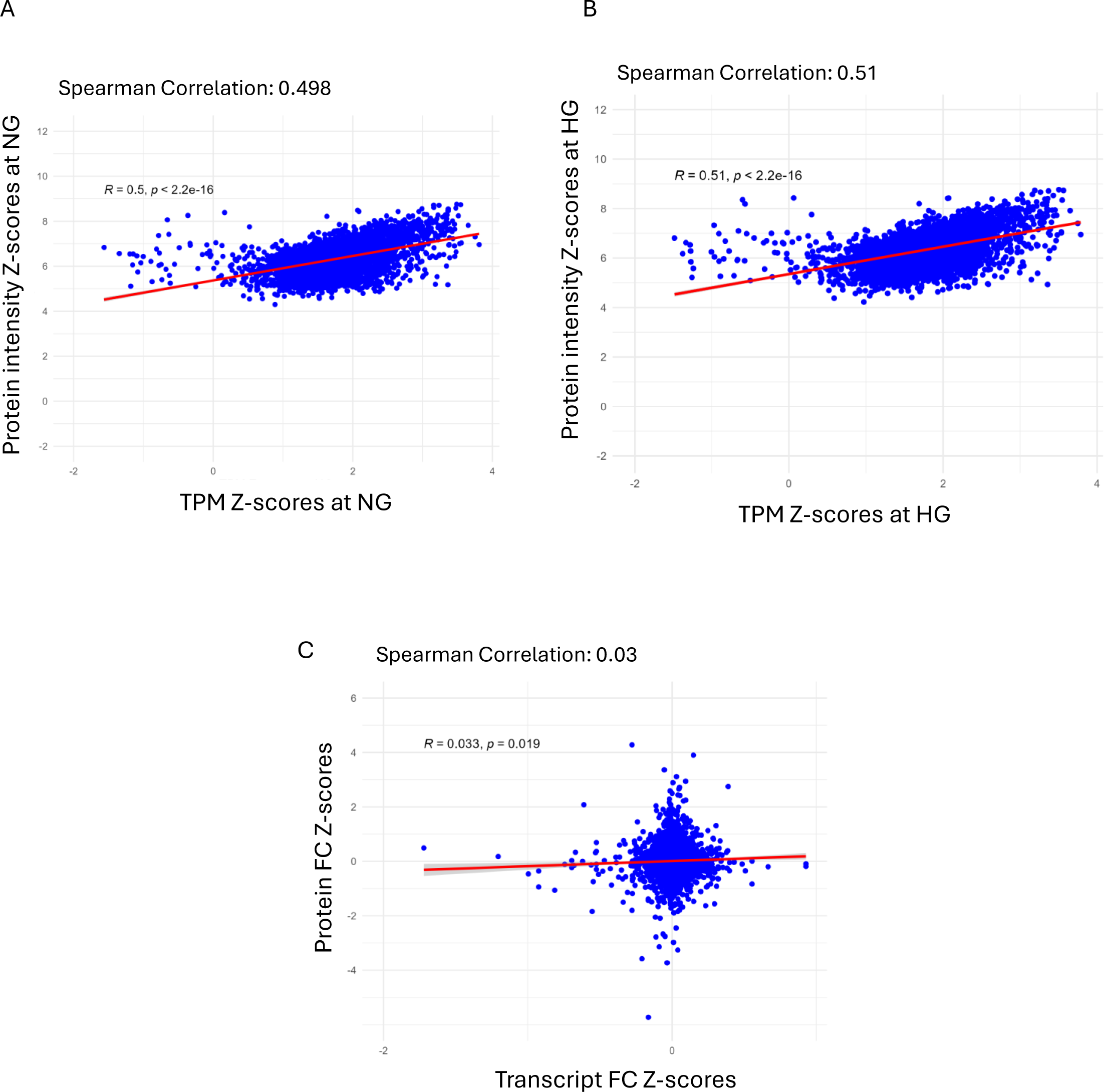
High glucose induces transcription-translation dissociation in MLO-Y4 cells. **(A)** Scatter plot represents spearman correlation of co-expressed transcripts and proteins under NG conditions and **(B)** HG conditions. **(C)** Spearman correlation of Log10 FC of xo-expressed transcripts and proteins.

### 3. Distinct RNA–protein profiles underly cellular response to HG stress

To investigate the biological functions associated with genes showing positive and negative expressions between the transcriptome and proteome, an enrichment analysis was performed using the DAVID platform. The analysis identified distinct enrichment patterns among the four expression groups, reflecting parallel or inverse expression between transcription and translation (Figure 4, Supplementary Table 7).

**Figure 4:**
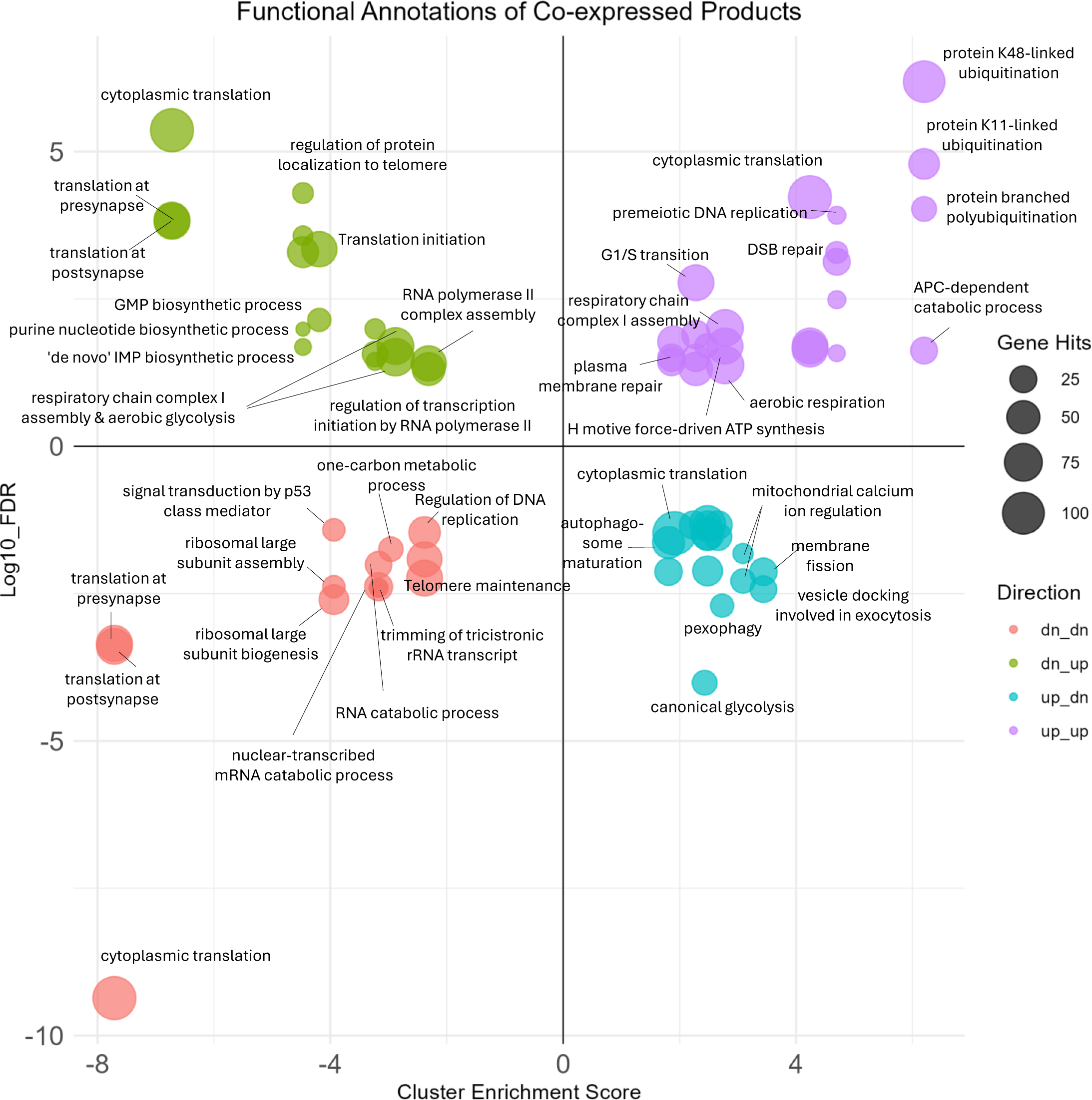
High glucose induces distinct RNA–protein relationships that define cellular response to stress. Up_up: upregulated at both RNA and protein. Dn_dn: downregulated at both RNA and protein. Up_dn: upregulated at RNA, downregulated at protein. Dn_up: Downregulated at RNA and upregulated at protein.

Genes upregulated at both mRNA and protein levels were enriched in pathways linked to protein ubiquitination and catabolic turnover, suggesting increased demand for proteostasis. High glucose induced higher enrichment of pathways associated with cell-cycle activity, and DNA replication and repair. Notably, the strongest enriched terms were K48- and K11-linked ubiquitination, branched polyubiquitin chains and anaphase promoting complex (APC) proteolysis consistent with cell cycle and stress-induced degradation of damaged or misfolded proteins [34], other enriched terms included regulation of translation, and oxphos.

Consistently downregulated genes were enriched in pathways associated with translation, ribosome assembly and biogenesis, RNA metabolism, and DNA maintenance. The top enriched terms were translation and ribosomal biogenesis. Additionally, RNA, mRNA catabolism, as well as tRNA surveillance signals were enriched. The downshift of ribosomal translation in addition to RNA degradation pathways points to a decrease in global translation capacity coupled to RNA quality control.

Transcriptionally activated programs that did not materialize at the protein level (up_dn) were top enriched in pathways associated with vesicle trafficking, mitochondrial calcium ion homeostasis. Enriched terms also included pexophagy or autophagy targeted at peroxisomes and maturation of autophagosomes, cytoskeletal reorganization pathways, cell cycle progression and glycolysis. On the other hand, the inverse pattern enrichment with proteins increased despite reduced transcripts were significantly enriched in cytoplasmic translation, telomeric protein recruitment, purine synthesis (IMP and GMP) and mitochondrial aerobic respiration.

Together, these four expression patterns reveal a multilayered regulatory landscape in which transcriptional and translational outputs diverge across distinct biological programs. Regulation of expression of RNAs and proteins highlighted a control of cell-cycle progression, proteostasis, RNA quality control translation capacity, and bioenergetic program activation. Translation-related themes appear in every pattern with varying degrees of enrichment suggesting nuanced shifts in translational activity. These contrasts underscore that transcriptional activation does not uniformly predict translational output and emphasize the presence of post-transcriptional and translational checkpoints shaping the cellular response. The functional implications of these relationships are addressed in the Discussion.

## Discussion

In this study, we examined how RNA–protein regulatory layers in osteocytic cells respond to hyperglycemic levels of glucose. By integrating RNA-sequencing with quantitative proteomics, we moved beyond single-layer analyses to assess how transcription and translation are coupled under basal conditions and, critically, how this relationship is remodeled under stress.

Transcriptomics analysis revealed that hyperosmotic stress from either high glucose or mannitol triggers a shared metabolic reprogramming in osteocytic cells, characterized by a shift from oxphos to aerobic glycolysis. These shared osmoadaptive responses involved the upregulation of key metabolic pathways, including lipid metabolism, the pentose phosphate pathway, and the TCA cycle [35]. This collective shift resembles a Warburg-like effect, where the cell moves toward aerobic glycolysis to enhance ATP generation and meet acute high energy demands characteristic of cellular stress [35]. Supporting this metabolic rewiring, is the upregulation of the Hif1ꭤ pathway, a known inducer of the oxphos-to-glycolysis switch [36]. Concurrently, this adaptation is accompanied by a transient suppression of protein translation pathways, indicated by the reduced enrichment of aminoacyl-tRNA biosynthesis.

When osmolarity was controlled for, distinct high glucose–specific transcriptional programs emerged. These were marked by a further suppression of oxidative metabolism alongside activation of selected anabolic and inflammatory pathways. High glucose triggered a coordinated downregulation of mitochondrial processes, including mitophagy; a key quality-control pathway for removing damaged mitochondria as well as antioxidant defense mechanisms [37,38].

Concurrently, HG specifically upregulated pathways related to the ECM and inflammatory responses in addition to key anabolic signaling axes, such as the PTH and Wnt-Lrp pathways, were upregulated, suggesting an initial push toward processes known to increase glucose uptake and glycolysis [39-42]. Interestingly, annotations related to bone resorption and rheumatoid arthritis were downregulated, suggesting a diminished capacity for osteocyte lacunar remodeling and recruitment of necessary cell pools for bone resorption.

Our integrative analysis at baseline culture conditions (normal glucose) showed a moderate correlation between transcript abundance and protein expression of co-expressed products. We also identified significant layer-specific patterns in our datasets: transcripts found exclusively at the RNA level were enriched for genes encoding membrane-associated or secreted proteins, which are often challenging to detect via mass spectrometry due to their physiochemical properties [28,30]. Conversely, proteins detected exclusively at the proteomic level were dominated by mitochondrial enzymes and cytoskeletal components with long protein half-lives, high stability, and slow turnover [31-33,43]. This highlights the osteocyte capacity in buffering the proteome, which can maintain basal levels of essential, long-lived proteins despite low acute transcriptional input of mitochondrial proteins and cytoskeletal mRNAs.

The central finding of this work is the identification of four discrete regulatory patterns that collectively define the multilayered osteocytic response to high glucose and reveal post-transcriptional mechanisms that modulate protein output relative to mRNA levels during metabolic stress.

The first pattern, upregulation at both RNA and protein levels, suggests increased demand for proteostasis elucidated by the enrichment of protein ubiquitination pathways. This group also showed upregulation of cell-cycle activity genes (like DNA synthesis and repair) and components of the electron transport chain (ETC), likely due to the overproduction of electron donors by the TCA cycle to complex I of ETC (NADH and FADH_2_) [44,45].

The second pattern, characterized by coordinated downregulation at both RNA and protein levels, was dominated by decreased translational activity, including reduced ribosomal biogenesis and diminished expression of rRNA, tRNA, and translation associated factors. This repression extended to one-carbon metabolism, a critical pathway linking core metabolism to the synthesis of nucleotides and amino acids [46]. This pattern is often seen in generalized stress responses where cells heavily manage energy resources [47].

The uncoupled patterns highlighted enrichment of critical translational and bioenergetic pathways in the cell. Transcriptionally activated but translationally suppressed genes, included pathways related to organelle autophagy, as well as upregulation of canonical glycolysis mRNAs hexokinase *(Hk1)* and proteoforms of phosphofructokinase *(Pfk)* unmatched at the protein level which are rate limiting steps in glycolysis. Whether this is true translational suppression can’t be verified from these data however, it is possible that increased mRNA levels is a transcriptional response to increased glycolytic overload in the face of high extracellular glucose. The inverse behavior were transcripts decreased while protein levels were maintained or increased was majorly enriched in factors involved in cytoplasmic translation, including eukaryotic initiation factors 4E and 4G (*Eif4e* and *Eif4g2*). These initiation factors are sub-stoichiometric components of the translational machinery, and their abundance is critical for setting baseline cap-dependent translational capacity whereas EIF4G2 is known to participate in cap-dependent translation in response to stress [48,49]. The enrichment of translation-related annotations across all four regulatory patterns indicates that high glucose reshapes translation by regulating the machinery responsible for mRNA transcription and protein synthesis. The data suggest a remodelling of various components and modulators of the translational machinery to regulate how osteocytes allocate their translational capacity under metabolic stress. Collectively, our data shows that osteocytes respond to stress in modular, multi-patterned way in which metabolic rewiring, proteostasis, and translational control are differentially engaged. A key next step is to define, which of these RNA–protein programs represent early adaptive responses that can be targeted in early hyperglycemic skeletal injury.

## Material and Methods

### Cell culture and experimental design

MLO-Y4 cells (AddexBio Technologies) were cultured at standard conditions (37 ֯Ϲ, 5% CO_2_) overnight in alpha-minimum essential medium (ꭤ-MEM; Wako Pure Chemical Industries, Osaka, Japan) containing 10% fetal bovine serum (FBS; Biowest, Nuaillé, France), 100IU/mL penicillin G, 100 μg/mL streptomycin (1% P/S) containing 5.5 mM glucose (standard medium). After that, the media was replaced with fresh 5.5mM glucose (normal glucose, NG), 25 mM glucose (Sigma-Aldrich, St. Louis, MO, USA) (high glucose, HG) containing media or 5.5 mM glucose supplemented with 19.5 mM mannitol (osmotic control, M) (Sigma-Aldrich) for 3 days.

### RNA sequencing

Total RNA was extracted using Qiagen RNeasy kit (QIAGEN, Hilden, Germany) with a DNase digestion step (QIAGEN) according to manufacturer’s instructions. The samples were processed by Azenta, Japan as follows: samples were screened and quantified on an Agilent 4200 Tapestation with RNA ScreenTapeR. The resultant RIN (RNA integrity number) was 9.9 or 10 for all samples. mRNA was enriched using NEBNext Poly(A) mRNA Magnetic Isolation Module (NEB) and RNA library preparation was done NEBNext Ultra II Directional RNA Library Prep Kit (Illumina E7760). Libraries were indexed, pooled, and subsequently run on an Illumina Novaseq to obtain strand-specific 150 bp paired-end and 50M reads depth. FASTQ files were then used for bioinformatics analysis. The quality of the raw sequencing data was checked by Azenta, Japan. The overall Mean Quality Score (Q Score) for the entire run was calculated to be 35.84. Furthermore, sequencing accuracy was confirmed by the high percentage of bases with a quality score of 30 or greater (Q30), with 93% of all bases across the run achieving this threshold. All sequencing experiments were done in triplicates.

### RNA sequencing analysis

Raw reads were processed to remove adapter sequences and low-quality bases using Trimmomatic. High-quality, filtered reads were aligned to the were aligned to the mouse reference genome (M35, GRCm39) using the aligner HISAT2. Gene and transcript counts were quantified using FeatureCounts with default parameters. Differential expression analysis was performed using were identified using Limma-Voom, applying TMM normalization. Genes with an adjusted p-value (FDR) of less than 0.05 and a log2-fold change greater than 1 were considered differentially expressed.

### Computational processing of transcriptomic data for integrative analysis

All transcriptomic analyses were performed in R, except for gene identifier conversion, which was carried out using the Ensembl BioMart web interface. TPM values were calculated by dividing raw read counts by transcript length in kilobases to obtain reads per kilobase (RPK), followed by scaling so that the sum of RPK values per sample = one million. Transcripts were filtered by expression level with transcripts TPM = 0 in any replica of 3 biological replicas filtered out. Gene identifiers were standardized using Ensembl release 115 with the GRCm39 mouse gene annotation. After identifier unification, the dataset was restricted to protein-coding genes to ensure compatibility with the proteomic dataset.

### Preparation of samples for mass spectrometry proteomics

5 samples per group were used for protein expression analysis. Samples were digested using the phase-transfer surfactant (PTS)-aided trypsin digestion protocol as described previously [50]. Briefly, the samples were suspended in 200 μL of PTS buffer (12 mM sodium deoxycholate, 12 mM sodium lauroyl sarcosinate in 100 mM Tris-HCl, pH 9.0) containing protease inhibitors. The samples were incubated on a heating block at 95 °C for 5 min and then sonicated for 20 min. The extracted proteins were quantified with a BCA protein assay kit and reduced with 10 mM DTT for 30 min at 37 °C, followed by alkylation with 50 mM 2-iodoacetamide (IAA) for 30 min at room temperature in the dark. The samples were diluted 5-fold with 50 mM ammonium bicarbonate. The proteins were digested with lysyl endopeptidase (LysC) and trypsin overnight at 37 °C in a shaking incubator. 1 mL of ethyl acetate was added to 1 mL of the digested solution, and the mixture was acidified with 0.5% trifluoroacetic acid (TFA) (final concentration). The samples were vortexed for 2 min and centrifuged at 15,800 × g for 2 min to separate the aqueous and organic phases. The aqueous phase was collected, dried, and the samples were resuspended in 1% TFA/5% acetonitrile (ACN) solution and loaded with a StageTip containing SDB-XC (upper) and SCX (bottom) Empore disk membranes [51,52]. Then they were eluted from the StageTip by 4% TFA with 30% ACN in 500 mM ammonium acetate and 30% ACN in 500 mM ammonium acetate. The sample was dried and resuspended in 0.5% TFA with 5% ACN. The peptide concentration was also measured on NanoDrop (Thermo Fisher Scientific) with an absorbance at 205 nm and an extinction coefficient of 31[53], and normalized to 80 ng/μL.

### Mass spectrometry proteomics

Mass spectrometry proteomics was carried out in the data-independent acquisition (DIA) mode with a FAIMS Pro Duo interface connected to an Orbitrap Fusion Lumos Tribrid mass spectrometer (MS). LC was performed on an Ultimate 3000 pump and an HTC-PAL autosampler using self-pulled needle columns (150 mm length, 100 μm ID, 6 μm needle opening) packed with Reprosil-Pur 120 C18-AQ 1.9 μm reversed-phase material [54]. The injection volume was 5 μL, and the flow rate was 500 nL/min. Separation was achieved by applying a three-step linear gradient of 4-8% ACN in 5 min, 5-32% ACN in 90 min, 32-80% ACN in 5 min, and 80% ACN for 10 min in 0.5% acetic acid. Data-independent acquisition (DIA) was performed on an Orbitrap Fusion Lumos Tribrid mass spectrometer (Thermo Fisher Scientific) equipped with a nano-electrospray ion source and a FAIMS Pro interface. The instrument was operated in positive ion mode with a spray voltage of 2.4 kV and an ion transfer tube temperature of 250 °C. FAIMS was run in standard resolution mode with a total carrier gas flow of 4 L/min, and compensation voltages (CVs) of −40 V and −60 V were applied in separate experiments. Full MS scans were acquired in the Orbitrap analyzer at a resolution of 60,000 (at m/z 200) over an m/z range of 400–950 using quadrupole isolation, with the AGC target set to the standard value (4 × 10^5^) and the maximum injection time controlled automatically. DIA MS/MS scans were acquired in the Orbitrap using higher-energy collisional dissociation (HCD) at a normalized collision energy of 30%, at a resolution of 15,000 (at m/z 200), with an AGC target of 1 × 10^6^ and a maximum injection time of 22 ms. For each FAIMS CV, 25 sequential quadrupole isolation windows with variable widths between 16 and 40 m/z were used to cover the precursor m/z range 400–950. Fragment ions were recorded in the Orbitrap over an m/z range of 200–2000, with the first mass of MS/MS scans set to m/z 200.

### DIA data processing and protein quantification

DIA raw files were processed using DIA-NN (version 2.2.0) [55]. A spectral library was generated from a library-free search workflow, using a false discovery rate (FDR) threshold of 1% at the peptide and protein levels. DIA data were searched against the [Mus musculus, UniProt, downloaded on 2025-01-16] with carbamidomethylation of cysteine as a fixed modification and methionine oxidation and protein N-terminal acetylation as variable modifications.

### Integrative analysis

After standardizing gene and protein identifiers and removing duplicates, transcript and protein abundance matrices were merged by unified gene symbol. Shared and layer-exclusive products were then filtered. For the baseline correlation analysis, mean TPM values (genes) and mean protein intensities (proteins) under normal glucose conditions were assessed using Spearman correlation. To examine how transcriptional and proteomic fold-changes relate under perturbation we computed log10 fold-change values for both RNA and protein levels to calculate Spearman correlation between transcript-level and protein-level responses as well as dividing responses into 4 groups based on their correlation coefficient. All calculations, transformations, and statistical analyses were performed in R.

### Functional enrichment analysis

Functional enrichment analysis was performed using the Database for Annotation, Visualization, and Integrated Discovery (DAVID) [27] or eVITTA’s pre-ranked gene set enrichment analysis (GSEA) [56]. The analysis included Gene Ontology (GO) biological process (BP) and cellular component (CC) categories using the Mus musculus background. Terms with a Benjamini-adjusted p-value (FDR) ≤ 0.05 and Cluster Enrichment Score (CES) ≥ 1.3 were considered for further interpretation in RNA-protein pattern analysis.

## Data availability

Raw sequencing data is deposited in Sequence Read Archive under the project ID PRJNA1371605. Data can be obtained from the following <u>link</u>.

The proteomics data have been deposited in the ProteomeXchange Consortium via the jPOST partner repository (https://jpostdb.org) [57] under the identifiers PXD071808 (ProteomeXchange) and JPST004230 (jPOST).

Any additional information required to reanalyze the data reported in this paper is available from the lead contact upon request.

## Acknowledgements

This work is supported by The FRIS Creative Interdisciplinary Collaboration Program and a Japan Society for the Promotion of Science, JSPS Kakenhi grant #25K20447 to AM.

---

**Supplementary Figure 1:** Volcano plot comparing the log2 FC and - log10 adjusted P value of RNA-seq data in MLO-Y4 cells comparing NG vs M.

**Supplementary Table 1:** GSEA comparing HG vs NG.

**Supplementary Table 2:** GSEA comparing M vs NG.

**Supplementary Table 3:** GSEA comparing HG vs M.

**Supplementary Table 4:** GSEA of shared pathways between HGvsNG and MvsNG.

**Supplementary Table 5:** Protein-coding transcripts identified in RNA-seq.

**Supplementary Table 6:** Co-expressed products at RNA and protein level.

**Supplementary Table 7:** Functional annotation of 4 Patterns of association between co-expressed RNAs and proteins.

