## Supplementary figures and images for "Rewiring of RNA–protein coupling in osteocytes in response to hyperglycemic levels of glucose"

### Supplementary Figure 1

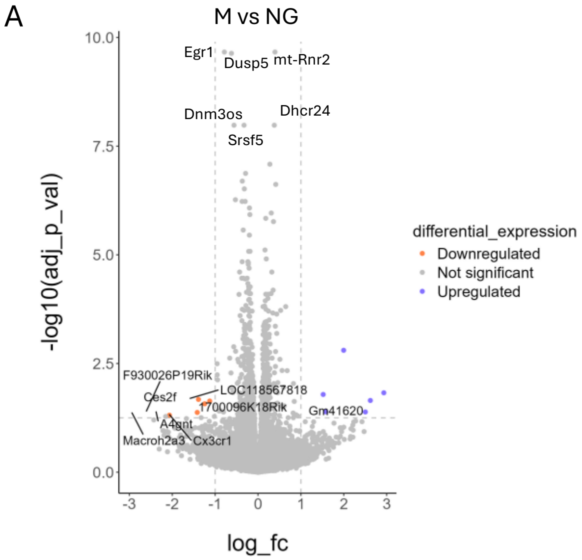
